# A sparse, spatially biased subtype of mature granule cell is preferentially recruited in hippocampal-associated behaviors

**DOI:** 10.1101/804393

**Authors:** Sarah R. Erwin, Weinan Sun, Monique Copeland, Sarah Lindo, Nelson Spruston, Mark S. Cembrowski

**Affiliations:** Dept. of Cellular and Physiological Sciences, Life Sciences Institute, University of British Columbia 2350 Health Sciences Boulevard, Vancouver, BC, Canada; Janelia Research Campus, Howard Hughes Medical Institute 19700 Helix Dr., Ashburn, VA 20147, USA; Djavad Mowafaghian Centre for Brain Health, University of British Columbia 2215 Wesbrook Mall, Vancouver, BC, Canada

## Abstract

Animals can store information about experiences by activating specific neuronal populations, and subsequent reactivation of these neural ensembles can lead to recall of salient experiences. In the hippocampus, granule cells of the dentate gyrus participate in such memory engrams; however, whether there is an underlying logic to granule cell participation has not been examined. Here, we found that a broad range of novel experiences preferentially activates granule cells of the suprapyramidal blade relative to the infrapyramidal blade. Motivated by this, we identified a suprapyramidal-blade-enriched population of granule cells with distinct spatial, morphological, physiological, and developmental properties. Via transcriptomics, we mapped these traits onto a sparse and discrete granule cell subtype that was recruited at a ten-fold greater frequency than expected by subtype prevalence, constituting the majority of all recruited granule cells. Thus, a rare and spatially localized granule cell subtype is intrinsically predisposed to activation during hippocampal memory formation.

## INTRODUCTION

The hippocampus is a brain region critical for episodic memory (Scoville and Milner, 1957), spatial navigation (O’Keefe and Nadel, 1978), emotion (Kjelstrup et al., 2002), and representation of a wide range of other internal and external states (Aronov et al., 2017; Ciocchi et al., 2015; Hitti and Siegelbaum, 2014; MacDonald et al., 2011). A central focus of hippocampal neuroscience lies in understanding the neurobiological substrates of this diverse functionality. One approach to identifying these substrates is centered around the perspective of cell types: identifying groups of cells that covary in specific properties, and via inference or further experimental assay, mapping such cell types onto functional contributions.

Classically, such cell-type divisions have been defined at a relatively broad level – for example, granule cells of the dentate gyrus, and pyramidal cells of regions CA3 and CA1 (Ramón y Cajal, 1911). More recently, an accumulation of evidence has emerged that such broadly defined hippocampal cell types can exhibit marked within-cell-type heterogeneity. Such work has largely focused on pyramidal neurons, wherein the long-range projections of these cells allow specific circuits to be experimentally manipulated and interpreted (Berns et al., 2018; Cembrowski et al., 2018a; Cembrowski and Spruston, 2019; Ciocchi et al., 2015; Jimenez et al., 2018; Okuyama et al., 2016; Soltesz and Losonczy, 2018; Spellman et al., 2015; Xu et al., 2016). In stark contrast to output pyramidal cells, markedly little is known about the subtype-specific decomposition of mature granule cells (GCs) of the dentate gyrus (DG), which form the local input layer of the hippocampus (Scharfman, 2007).

The relative lack of subtype-specific insight into mature GC organization and recruitment is striking, as GCs are a focal point of engram memory research (Bernier et al., 2017; Chen et al., 2019; Denny et al., 2014; Guskjolen et al., 2018; Kheirbek et al., 2013; Liu et al., 2012; Park et al., 2016; Ramirez et al., 2013; Redondo et al., 2014; Ryan et al., 2015; Tonegawa et al., 2015). Notably, previous work potentially hints at the existence of DG subtype-specific recruitment: interventional access-and-manipulate approaches anecdotally exhibit GCs in the suprapyramidal blade preferentially recruited relative to the infrapyramidal blade (e.g., Chen et al., 2019; Liu et al., 2012; Ramirez et al., 2013; Redondo et al., 2014; Ryan et al., 2015). This across-blade difference has been more formally examined in observational immediate-early gene experiments, wherein activity has been shown to biased to the suprapyramidal blade in novel environment exposure (Chawla et al., 2005; Chawla et al., 2018; Guenthner et al., 2013; Penke et al., 2011; Ramirez-Amaya et al., 2005). To date, the degree to which this blade-specific recruitment generalizes across behaviors, maps onto GC subtypes, and ultimately relates to function are unknown.

Here, we sought to understand whether functional recruitment of GCs could be interpreted and predicted according to underlying subtype-specific rules. Beginning with activity labeling, we found blade-specific activity differences emerged across a wide range of disparate behavioral paradigms. We registered this functional heterogeneity with underlying differences in GC morphology, physiology, spatial location, and gene expression, and mapped this multimodal heterogeneity onto a discretely separate and rare mature GC subtype. Finally, we show that this subtype is preferentially activated by experiences associated with hippocampal-dependent memory, accounting for 70-80% of recruited GCs despite comprising only 5% of the total GC population. This work leads to the unexpected conclusion that subtype-specific heterogeneity exists and predicts recruitment at the first stage of hippocampal processing.

## RESULTS

### Behavior preferentially tags suprapyramidal blade granule cells

We began by confirming that GCs of the suprapyramidal blade of the DG (i.e., located proximal to stratum lacunosum-moleculare; Fig. 1A) were selectively activated by exposure to a novel environment, as previously shown (Chawla et al., 2005; Chawla et al., 2018; Guenthner et al., 2013; Penke et al., 2011). To do this, we introduced double-transgenic FosTRAP (i.e., cFos-cre(ERT2)) x Ai14 (i.e., LSL-tdTomato) mice (Guenthner et al., 2013) to a novel environment and, after 20 minutes of exploration, administered 4-OHT prior to returning animals to their homecage (Fig. 1B). This inducible transgenic system enabled fluorescent tagging of cells active during novel environment exploration, and labeled cells that were largely restricted to the suprapyramidal blade (Fig. 1C).

**Fig. 1.**
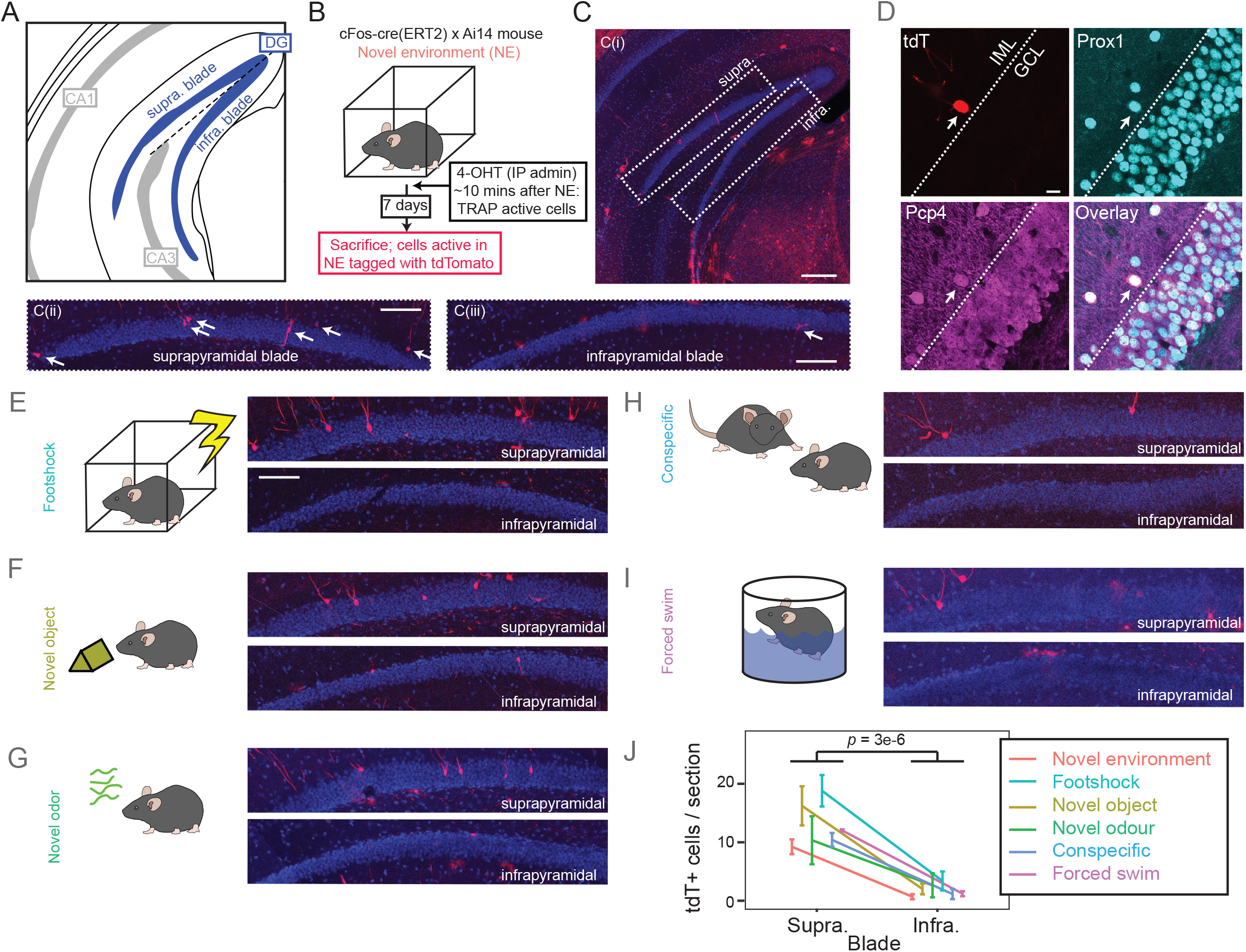
Granule cells of the suprapyramidal blade of the dentate gyrus are preferentially activated across a range of behavioral paradigms. A. Atlas illustrating the blades of the DG in a coronal section. Modified from (Paxinos and Franklin, 2004). B. Schematic of tdTomato labeling of cells active during novel environment exploration. C. Optical section of tdTomato labeled cells following NE exploration. C(i) depicts overview, whereas C(ii) and C(iii) depict expansions of the suprapyramidal and infrapyramidal blades, respectively. Arrows denote labeled cell bodies with morphologies consistent with GCs. Scale bars: 200, 100, and 100 µm, respectively. D. Labeled cells were positive the DG GC markers Prox1 and Pcp4, as assayed via immunohistochemistry. Dashed line denotes boundary between inner molecular layer (IML) and granule cell layer (GCL). Scale bar: 5 µm. E-I. As in C(ii,iii), but for a variety of behavioral paradigms. Scale bar: 100 µm. J. Summary of the number of labeled cells, per 100 µm-thick section, for the two blades of the dentate gyrus. Central tendency and error bars denote mean ± SEM.

In general, tdTomato-labeled cells exhibited morphological properties consistent with excitatory granule cells. For example, labeled cells exhibited apical dendritic trees (Fig. 1C), and also showed prominent dendritic spines (Supplemental Fig. 1). Remarkably, many tdTomato-labeled cells also exhibited a cell body at or beyond the interface between the granule cell layer and the inner molecular layer (IML) (e.g., Fig. 1D and Supplemental Fig. 1; 43% of examined neurons had cell bodies clearly displaced in or beyond the IML, n = 56/130). Despite these cell bodies being displaced relative to the classical granule cell layer, multiple GC markers (Prox1 and Pcp4: Cembrowski et al., 2016b) labeled these cells (Fig. 1D; 100% of examined displaced tdTomato-labeled cells exhibited double labeling for both Prox1 and Pcp4; n=10/10).

We next examined whether this blade specificity generalized to other hippocampal-associated behavioral paradigms. Similar suprapyramidal blade-specific recruitment was seen following foot shock in a novel environment, introduction of novel objects or odors into the homecage, introduction of a conspecific, and participation in a forced swim test (representative images: Fig. 1E-I; summary data: Fig. 1J). In total, preferential suprapyramidal blade activation occurred in behaviors evoking memory, spatial navigation, socialization, and stress.

### Tagging is consistent with bona fide activity differences

We next performed control experiments to help interpret these previous results. First, we performed negative controls to examine the extent of activity labeling in other settings. Here, we found behaviorally induced labeling was much greater than in saline-control animals and animals receiving 4-OHT in their homecage (see also Guenthner et al., 2013), as well as animals transferred and handled in behavior room without a subsequent behavioral assay (Supplementary Fig. 2A-E). Next, we performed positive controls to ensure our results generalized across activity detection paradigms. In these experiments, preferential suprapyramidal blade activation was seen following endogenous Fos labeling (Supplementary Fig. 2F), as well as using other IEG targets (e.g., *Arc*: Supplementary Fig. 2G) (see also Chawla et al., 2005; Chawla et al., 2018; Penke et al., 2011; Ramirez-Amaya et al., 2005). Finally, we ensured that infrapyramidal blade GCs could be induced to express Fos by exogenous stimulation (Supplementary Fig. 2H) (see also Chawla et al., 2005). In collection, these controls suggest *bona fide* activity differences underlie suprapyramidal-blade-enriched behavioral labeling (Fig. 1).

### The suprapyramidal blade preferentially exhibits displaced granule cells

As our activity-recruited cells were both enriched in the suprapyramidal blade and exhibited cell bodies “displaced” into the ML (e.g., Fig. 1D), we hypothesized that there was an inherent blade-specific difference in displaced GCs. Consistent with this, *in situ* hybridization (ISH) and immunohistochemical (IHC) labeling of GCs revealed that vast majority of displaced GCs were found in the suprapyramidal blade (ISH: Fig. 2A, IHC: Fig. 2B; n=114/127, 89% and n=116/132, 90%, of displaced cells were associated with the suprapyramidal blade with ISH and IHC respectively). Similar results could be seen for other markers of GCs (e.g., *Prox1* and *Slc17a7*, Supplementary Fig. 3A,B). This revealed a previously unidentified and prominent difference in the blades of the DG under naïve, physiological conditions.

**Fig. 2.**
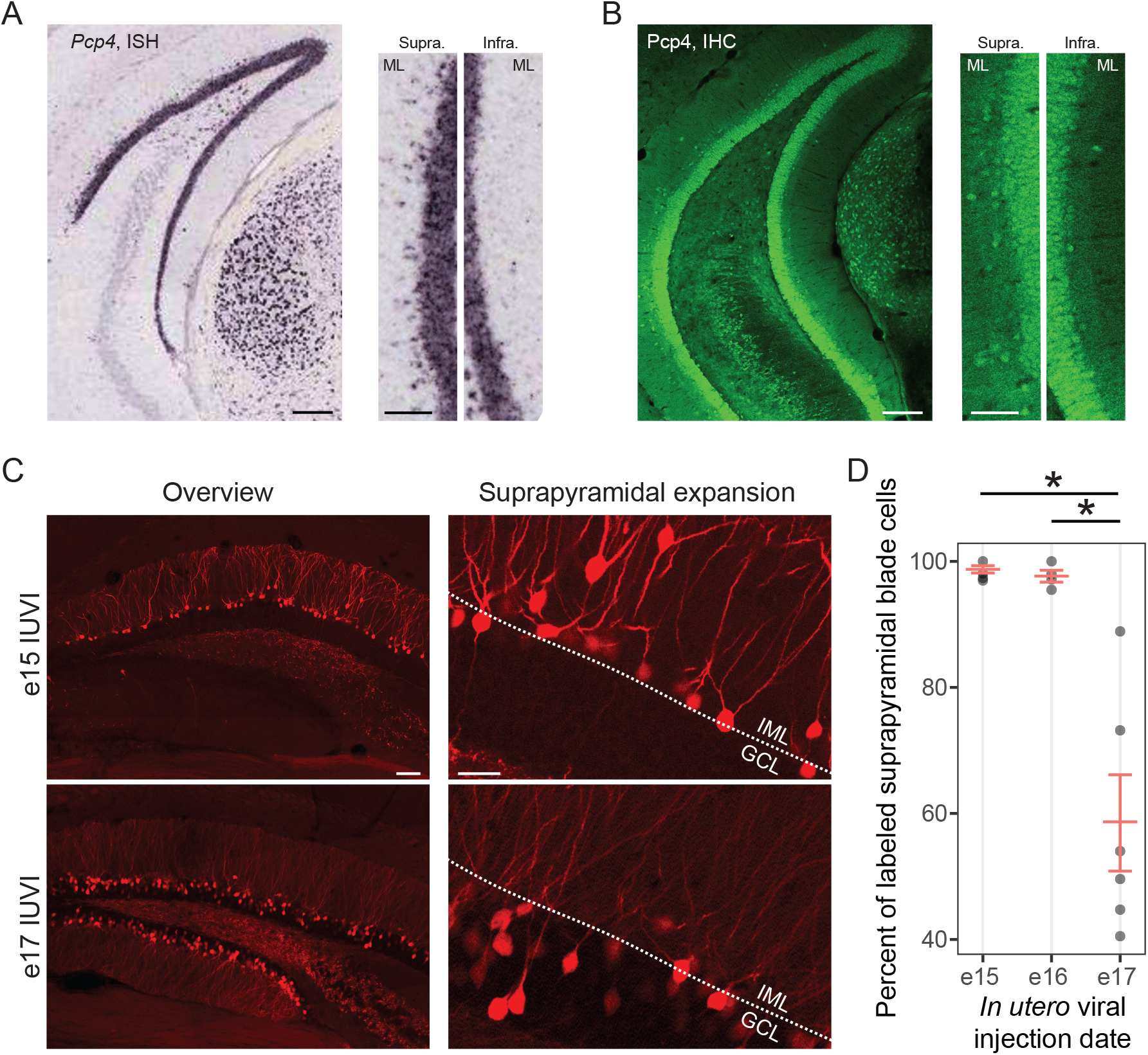
The blades of the dentate gyrus exhibit mature and birthdate differences. A. Left: ISH for the DG marker gene *Pcp4*. Scale bar: 200 µm. Right: magnification of the suprapyramidal and infrapyramidal blades. Note the suprapyramidal blade is prominently enriched for *Pcp4*-expressing cell bodies displaced into the MLs. Scale bar: 100 µm. B. As in (A), but for IHC detection of Pcp4 protein. C. Top row: Labeled granule cells in mature mice following birthdate labeling of neurons using *in utero* viral injections (IUVIs) at e15, shown in both overview (left) and expansion of the suprapyramidal blade (right). Scale bars, left and right: 100 µm and 25 µm. Bottom row: as in top row, but for *in utero* viral injections at e17. D. Summary of labeled cells across e15, e16, and e17 injection time points. Individual data points represent results from individual animals, and red lines with error bars reflect mean ± SEM for each time point.

Given the pronounced differences in the number of displaced GC between the two blades, in conjunction with the fact that GCs are born in a deep-to-superficial fashion (Angevine, 1965; Save et al., 2019), we further hypothesized that such differences may have a developmental origin. To resolve this, we performed birthdate labeling of DGs, injecting AAV2-CAG-FLEX-tdTomato into the GC-selective Rbp4-cre line (Cembrowski et al., 2016b) at different time points *in utero*. Labeled neurons following injections at e15 and e16 primarily exhibited properties of the “displaced” GC population: such cells exhibited atypical morphologies, with cell bodies bordering or within the molecular layers (Fig. 2C; cf. Fig. 1D). In stark contrast to this, labeled neurons following e17 injections were much more uniformly distributed across blades, and exhibited much less cell-body displacement outside of the GCL (Fig. 2C,D). In total, this illustrated a developmental origin consistent with the blade-specific GC displacement in maturity.

### Activity-labeled granule cells are consistent with semilunar granule cells

The displaced cell body location and broader dendritic branching of activity-labeled GCs resemble features of so-called semilunar granule cells (SLGCs) (Williams et al., 2007). In addition to these anatomical/morphological properties, SLGCs also have a markedly lower input resistance relative to classical GCs (Williams et al., 2007). Motivated by this previous work, we next used *ex vivo* morphological and electrophysiological techniques to identify the relationship between activity-labeled cells (defined by tdTomato expression) and SLGCs (previously defined according to morphological and electrophysiological criteria: Gupta et al., 2012; Larimer and Strowbridge, 2008, 2010; Save et al., 2019; Williams et al., 2007).

In mice that were exposed to a novel environment and activity tagged (as in Fig. 1A-D), we performed whole-cell recordings from both tdTomato-negative and tdTomato-positive cells in *ex vivo* brain slices (Fig. 3A,B). Recorded cells were filled with biocytin, allowing *post hoc* morphological reconstruction and analysis. Notably, tdTomato-positive cells typically exhibited differences in morphological and electrophysiological properties relative to tdTomato-negative cells (e.g., dendritic span, Fig. 3C; input resistance, Fig. 3D; see Table S1 for full summary of measured parameters and statistical tests). All of these features recapitulated previously described differences between SLGCs and classical granule cells (Larimer and Strowbridge, 2008, 2010; Williams et al., 2007), with tdTomato-positive cells corresponding to SLGCs in particular.

**Fig. 3.**
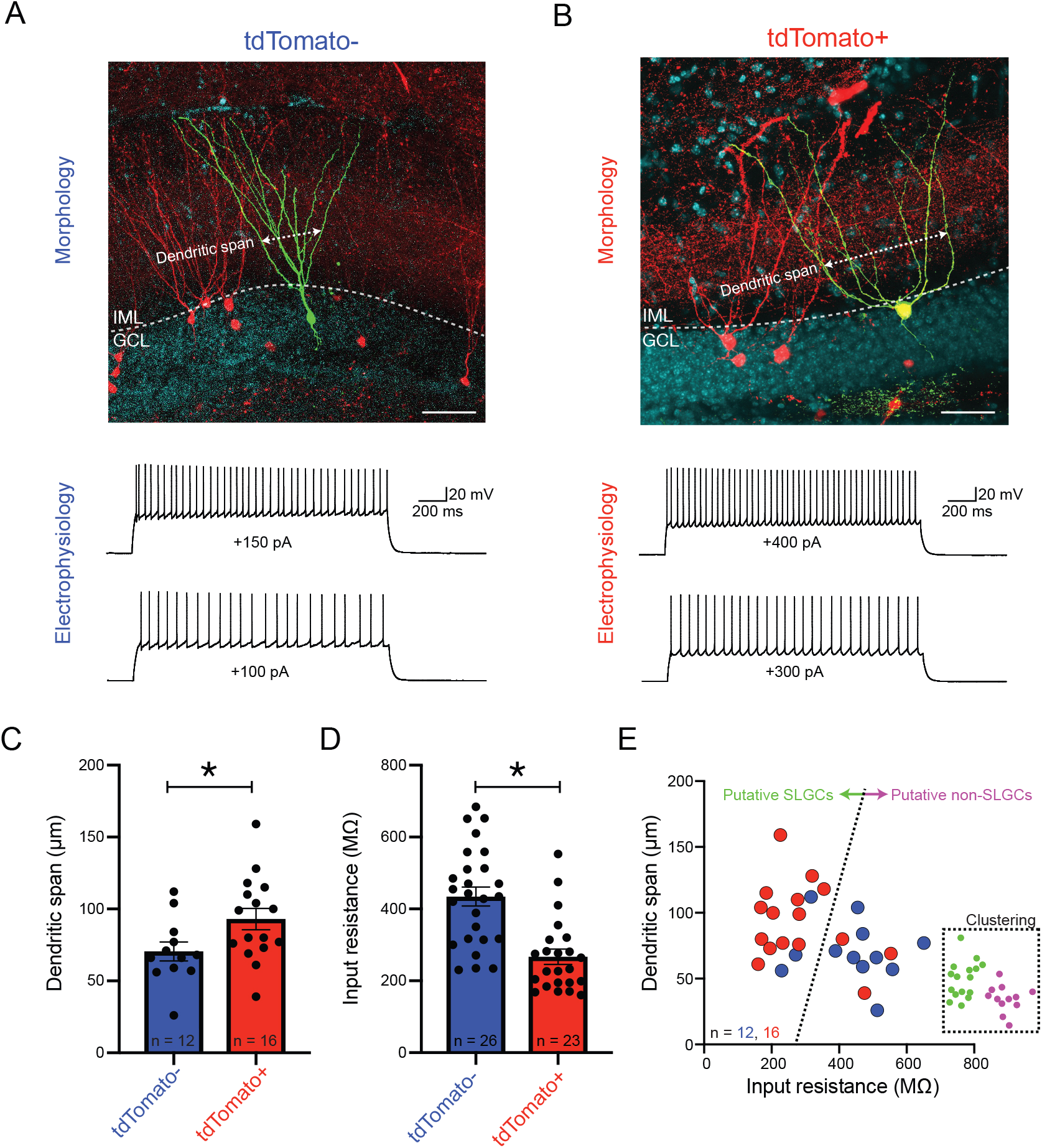
Activity-labeled neurons are consistent with semilunar granule cells. A. Top: Example morphology of a tdTomato-negative granule cell, recorded via whole-cell patch clamp *ex vivo*. Dendritic span, defined as the distance between the two outermost dendrites at 50 µm above the initial bifurcation of primary dendrite, is shown. Scale bar: 50 µm. Bottom: Example voltage responses following current step injection for the same cell. B. As in (A), but for a tdTomato-positive granule cell. C,D. Example of morphological (dendritic span, C) and electrophysiological (input resistance, D) properties of tdTomato-negative and tdTomato-positive cells. See Table S1 for full list of measured parameters and statistical tests. E. Two-parameter scatterplot of input resistance and dendritic span, illustrating separation that is recapitulated by k-means clustering (green and magenta points, inset).

Note that a perfect one-to-one correspondence between activity-labeled cells and SLGCs would not be expected: it is likely that some SLGCs are not recruited by novel environment exploration, whereas some classical GCs are recruited by this exploration. To investigate this, we performed a cluster analysis on our dataset. We found clusters largely agreed with activity labeling, and in particular suggested that putative SLGCs represented ~80% of the recruited cells (n=13/16 of tdTomato-expressing cells corresponded to one cluster: Fig. 3E).

### A distinct, discrete Penk-expressing subtype in the suprapyramidal blade

Motivated by activity labeling potentially having a subtype-specific basis, we next sought to identify whether GC heterogeneity adhered to a continuum or reflected discretely separated subclasses (Cembrowski and Menon, 2018). To do this in a quantitatively rigorous fashion, we analyzed a previously published single-cell RNA-seq (scRNA-seq) dataset that included 498 cells from the dentate gyrus (Habib et al., 2016). Combining nonlinear t-SNE visualization with graph-based clustering, we identified seven distinct DG cell classes (Fig. 4A). From these seven classes, four expressed marker genes associated with DG GCs (e.g., excitatory cell marker *Slc17a7* and GC marker *Prox1*, Fig. 4B-D; see also Supplementary Fig. 3C-H).

**Fig. 4.**
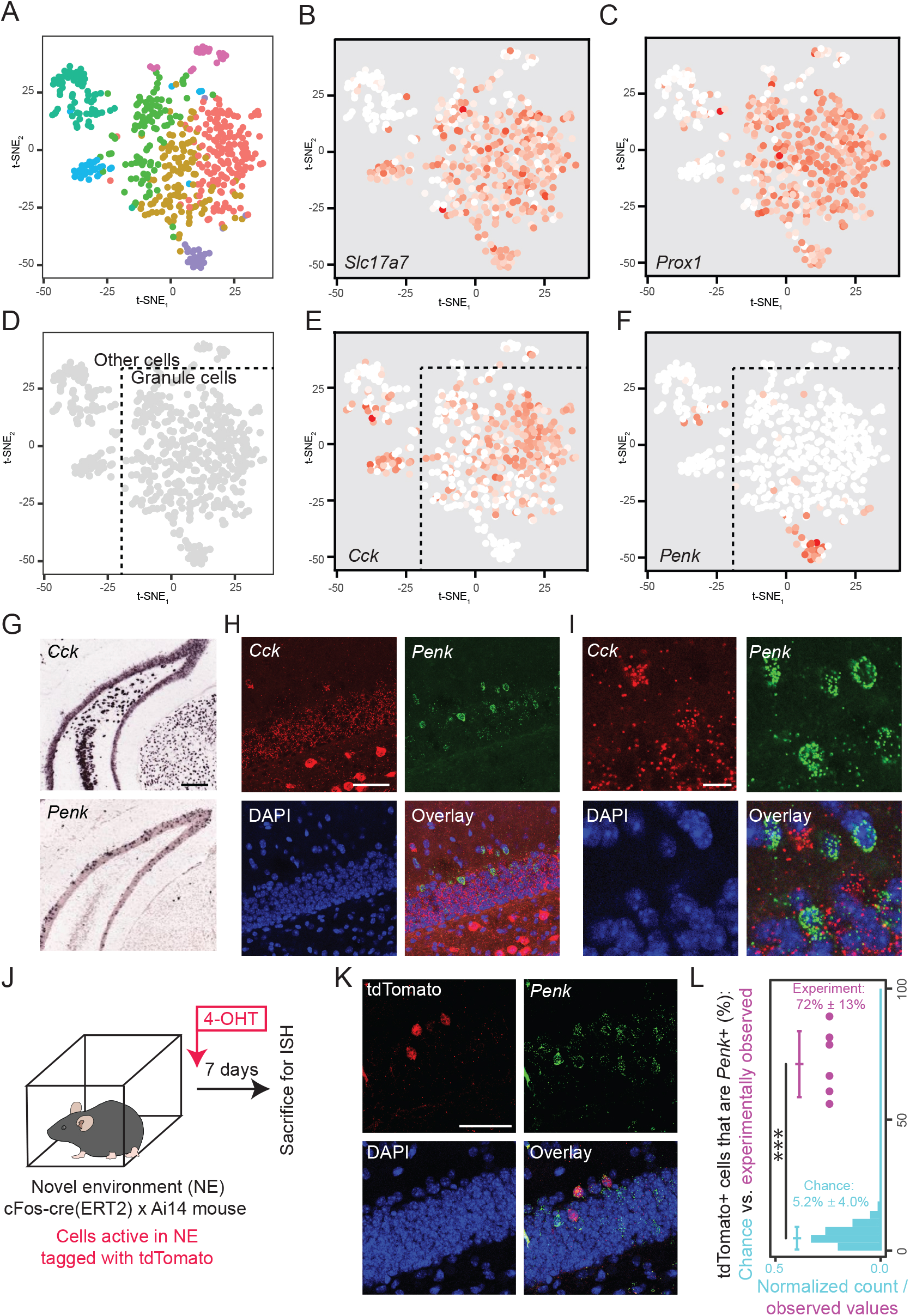
The DG embeds a sparse, blade-enriched discrete GC subtype that is preferentially recruited during behavior. A. tSNE visualization of scRNA-seq transcriptomes from the dentate gyrus. Colors denote different clusters of cells. B. Expression of *Slc17a7*, a marker of excitatory neurons. C. Expression of *Prox1*, a marker of GCs. D. Deconstruction of the scRNA-seq landscape into GCs and other cell types. E,F. Within the GC population, two subtypes of GCs can be identified based upon mutually exclusive expression of *Cck* and *Penk*. G. Single-color ISH of *Cck* and *Penk.* Scale bar: 100 µm. H,I. Overview (H) and magnification (I) of two-color smFISH of *Cck* and *Penk.* Scale bars: 50 µm and 10 µm. J. Illustration of behavioral paradigm to compare activity-labeled cells with *Penk*-expressing cells. K. Representative images of overlap between activity-labeled cells and *Penk-*expressing cells. Scale bar: 50 µm. L. Quantification of activity-labeled cells that also express *Penk*.

Within the GC dataset, a small and putatively rare *Penk*-expressing population was discretely separated from a broader collection of *Cck*-expressing cells (Fig. 4E,F). In addition to differences in these peptidergic markers, these populations also varied by a host of other functionally relevant genes, including those that regulate axon guidance and cell adhesion (*Slit1, Col6a1*), G protein signaling (*Rgs4*), cytoskeletal properties (*Nefm*), calcium binding (*Necab3*), and voltage-gated channels (*Scn3b*) (Supplementary Fig. 3I). Thus, it is likely that these two putative subtypes would vary in a host of higher-order properties, consistent with our previous results (Figs. 1-3). Notably, these differences were not explained by variation in the dorsal-ventral axis (Supplementary Fig. 3J,K), nor by maturity state (Supplementary Fig. 3L, in agreement with Fig. 2C), suggesting that a different feature might covary with these gene-expression differences.

Consequently, we examined whether the rare *Penk*-expressing population might correspond to the blade-enriched GC population. Remarkably, ISH labeling of *Penk*-expressing cells was found to be sparse, biased to cells near or within the molecular layers, and enriched in the suprapyramidal blade (Fig. 4G; 79% of labeled cells in suprapyramidal blade; n = 457/579 cells). Using two-color single-molecule fluorescent ISH (smFISH), we confirmed that *Penk* and *Cck* labeled nonoverlapping populations, in agreement with these peptidergic markers labeling discrete subtypes of GCs (92.5% of labeled cells exhibited mutually exclusive expression of either *Penk* or *Cck*, n = 414/458 counted cells). In particular, the *Penk-*expressing population labeled a rare subtype of GC, as expected from scRNA-seq (Fig. 4H,I; *Penk*-expressing neurons represented ~4.6% of all GCs, n=69/1498 counted cells; cf. 6.6% of all GCs in scRNA-seq dataset, n=25/380 in Fig. 4F).

### Penk-expressing granule cells are selectively recruited in behavior

Given the suprapyramidal blade enrichment and ML-displacement of *Penk*-expressing GCs, we sought to investigate whether this subclass preferentially participated in hippocampal-associated behavior (Fig. 1). As with previous behavioral experiments (Fig. 1B), cells were permanently labeled in response to novel environment exposure (Fig. 4J), with animals later sacrificed for smFISH subtype-specific identification. Remarkably, *Penk*-expressing cells exhibited a much higher propensity to be incorporated into active ensembles (72 ± 13% of activity-labeled cells exhibited *Penk* expression in six animals, mean ± SD, cf. 5.2 ± 4.0% expected by chance, *p* = 1.1e-5, Fig. 4K,L). Thus, cells activated by novel environment exploration largely conformed to a subclass of GCs expressing *Penk*, exhibiting a recruitment rate similar to SLGCs (cf. ~80%, Fig. 3E) and at an order of magnitude greater than predicted by prevalence alone (cf. *Penk*-expressing GCs comprising ~5% of all mature GCs, Fig. 4D,H).

## DISCUSSION

Although GCs are frequently examined as a model system for understanding the cellular underpinnings of memory, whether GC function can be interpreted and predicted according to GC subtypes remains uncertain. Our work here illustrates that the DG contains a pre-existing GC subtype that is suprapyramidal-blade-enriched and preferentially recruited during hippocampal-associated behavior (Figs. 1, 2). Such blade-enriched recruited cells display hallmarks of the atypical semilunar GCs, including a distinct morphology, cell body location, and electrophysiology (Fig. 3). This subtype is discretely separable from classical mature GCs according gene expression, and although quantitatively rare relative to classical GCs, constitutes the majority of all GCs recruited during behavior (Fig. 4). In total, our work here provides unexpected subtype-specific underpinnings of GC structural and functional variability, and will help to guide and interpret future experiments on the cellular basis of memory.

### Subtype-specific interpretation of non-uniform granule cell recruitment

A hallmark of granule cells is their low level of activity for a given environment (GoodSmith et al., 2017; Jung and McNaughton, 1993; Neunuebel and Knierim, 2012; Senzai and Buzsaki, 2017; Skaggs et al., 1996). When the same neurons are studied across environments, a small subset of granule cells accounts for most activity (Mizuseki and Buzsaki, 2013), with the mechanisms underlying this functional selectivity being unknown. Our findings here suggest this can be accounted for by GC subtype-specific contributions: the *Penk*-expressing GC subtype identified here accounts for 5% of the total GC population but for ~70-80% of the recruited GCs (Figs. 3, 4). In total, this amounts to *Penk*-expressing GCs being recruited at a frequency more than ten-fold predicted by prevalence alone, and comprising the majority of all recruited GCs.

The properties of this subtype-privileged recruitment are consistent with multiple other bodies of work. Recent correlative electrophysiology-morphology, combining juxtacellular recordings with morphological reconstructions of GCs, revealed that active GCs exhibit more complex dendritic arbors (Diamantaki et al., 2016) (see also Claiborne et al., 1990). In a complementary study using RNA-seq, *Penk* expression was found to be enriched in DG engram cells (Rao-Ruiz et al., 2019). These findings agree with the activity-biased DG subtype identified, which are characterized by morphologically complex dendritic arbors (Fig. 3) and *Penk* marker-gene expression (Fig. 4). In total, the subtype-specific organization uncovered here helps to provide a provide a framework to interpret these previous recruitment results.

The finding that a relatively rare subtype of GC dominates DG activity, while consistent with previous literature (Jung and McNaughton, 1993; Mizuseki and Buzsaki, 2013; Neunuebel and Knierim, 2012; Skaggs et al., 1996), is enigmatic. At first pass, this might suggest the overall computational capabilities of the dentate gyrus are much more limited than a raw count of GC would predict. However, such GC activity disparities may underscore a functional role: the *Penk-*expressing granule cells identified here may be sufficient to convey rapid and coarse features of the environment, whereas sparsely active classical granule cells may convey more nuanced information (Buzsaki and Mizuseki, 2014). In this way, the intrinsic architecture of the DG may support multiscale operations that provide computational and behavioral flexibility.

### Discovery and implications of blade-specific heterogeneity

Previous functional work, assaying activity via IEG labeling, has shown that the suprapyramidal blade is preferentially recruited during novel environment exploration (Chawla et al., 2005; Chawla et al., 2018; Guenthner et al., 2013; Penke et al., 2011). Such phenomenology could emerge from underlying cell-intrinsic and/or circuit differences between the blades of the DG. Our work here, revealing a preexisting GC subtype that is predisposed to recruitment, provides evidence for a cell-intrinsic mechanism. That such blade-specific activity is intrinsic, rather than reflecting long-range circuit inputs, is also in keeping with the lack of projection differences between the blades (Scharfman, 2007; van Groen et al., 2003; but see Wyss et al., 1979). It is important to note that local microcircuitry contributions, such as local excitatory IML recurrent connections from semilunar axon collaterals (Williams et al., 2007) or local inhibitory interneurons (Seress and Pokorny, 1981), may also augment and further amplify such cell-intrinsic differences.

Such blade-specific heterogeneity has important methodological implications for assaying GC activity going forward. With *in vivo* imaging technology becoming increasing available in neuroscience (Cai et al., 2016), it is critical to note that typically such technologies are used to interrogate only the suprapyramidal blade, and therefore likely provide a subtype-enriched interpretation of GC dynamics. Complementary techniques that allow concurrent activity readouts of both blades, such as IEG labeling or multisite electrophysiology, can circumvent this issue and will allow a more generalizable understanding of GC activity.

In addition to illustrating the existence of this across-blade heterogeneity, our results also provide a means of assaying the corresponding subtype-specific functional relevance. The marker genes identified here, *Penk* and *Cck*, provides a histological means of tagging non-canonical and canonical GCs for subtype-specific interpretation (e.g., Fig. 4). Complementing these observational experiments, transgenic animals that leverage the specific expression of these genes will enable interventional experiments and inference of causal relationships (Daigle et al., 2018; Harris et al., 2014; Taniguchi et al., 2011). Thus, the results here open multiple avenues for unraveling the subtype-specific rules of hippocampal-dependent memory and function.

---

**Supplemental Fig. 1. Representative depiction of activity-labeled cells and dendrites**.

Related to Figure 1.

Top: maximum intensity projection giving overview of activity-labeled cells in the suprapyramidal blade. Note displaced cell bodies and the presence of dendritic spines. Middle: expansion on a labeled cell. Bottom: expansion on labeled dendrites. Scale bars: 100, 10, and 20 µm, respectively.

**Supplemental Fig. 2. Control activity-labeling experiments.** Related to Figure 1.

A. Representative image showing absence of labeling following vehicle injections. Scale bar: 500 µm. B. Representative image showing activity-tagged cells for an animal in homecage. C. Representative image showing activity-tagged cells for an animal transported to the behavior room and handled, but not put through a behavioral paradigm. Arrows indicate activity-tagged (tdTomato-expressing) GCs in (A-C). D. Summary of negative control data (n=2 mice for vehicle injections, n=3 mice for each of homecage and behavior room transfer). E. For comparison to (D), summary of animals put through behavioral paradigms (as in Fig. 1J). Individual lines reflect individual animals, lines with error bars reflect pooled mean ± SEM, and statistical comparisons represent Mann-Whitney U-tests. F. Representative image showing preferential suprapyramidal labeling following novel environment exploration, as assayed through cFos IHC. Arrows indicate cFos+ cells in the granule cell layer and molecular layers. Note in general, the suprapyramidal was associated with 80% of cFos-labeled cells (n = 80/100 cells, 2 animals and 2 sections per animal). Scale bar: 200 µm. G. Representative image showing preferential suprapyramidal blade labeling following novel environment exploration, as assayed through *Arc* ISH. Arrows indicate *Arc-*expressing cells in the granule cell layer and molecular layers. Note that, in general, the suprapyramidal was associated with 70% of *Arc*-labeled cells (n = 276/395 cells, 2 animals and 6 sections per animal). H. Upper left: DAPI image of the infrapyramidal blade of the dentate gyrus. Upper right: mCherry expression, corresponding to injection of AAV2-hSyn-DIO-hM3D(Gq)-mCherry into the DG of Rbp4-cre mice. Lower left: immunohistochemical staining of cFos. Similar broad staining of the outer blade of the dentate gyrus was seen in 2 other animals. Lower right: overlay of all panels. Scale bar: 100 µm.

**Supplemental Fig. 3. Gene expression in the dentate gyrus.** Related to Figures 2 and 4.

A. Left: overview of *Prox1* expression in the dentate gyrus, a marker of granule cells. Scale bar: 500 µm. Right: expansion of the suprapyramidal (top) and infrapyramidal (bottom) blades. Scale bar: 100 µm. Note suprapyramidal blade enrichment of displaced granule cells. B. As in (A), but for the excitatory neuron marker gene *Slc17a7.* C. t-SNE visualization of single-cell transcriptomes, with coloring denoting cluster identity. D-G. Expression of the excitatory neuron marker *Slc17a7* (D), the DG GC marker *Prox1* (E), the inhibitory interneuron marker *Gad2* (F) and the glial marker *Slc1a2* (G). H. Cell-type-specific labeling of DG cell transcriptomes. I. Expression of subtype-enriched marker genes with neuronally relevant functional correlates. J,K,L. Expression of the dorsal DG marker gene *Lct* (J), the ventral marker gene *Trhr* (K), and the immature granule cell marker *Dcx* (L). Note that expression of each marker gene in (J,K,L) is dispersed across the GC clusters rather than exhibiting structured expression.

**Supplemental Table 1.** Morphological and electrophysiological properties of active and inactive granule cells. Related to Figure 3.

| | Resting membrane potential (mV) | Input resistance (MΩ) | AP amplitude (mV) | AP half width (ms) | AP threshold (mV) | Adaptation ratio | AHP amplitude (mV) | Dendritic span (μm) |
| --- | --- | --- | --- | --- | --- | --- | --- | --- |
| tdTomato- | -79.0 ± 0.9 | 434 ± 26 | 112.8 ± 2.2 | 0.993 ± 0.030 | -35.9 ± 0.7 | 0.30 ± 0.04 | 12.7 ± 1.0 | 71 ± 7 |
| tdTomato+ | -78.8 ± 1.0 | 267 ± 21 | 113.3 ± 3.6 | 0.972 ± 0.050 | -37.9 ± 1.8 | 0.46 ± 0.09 | 14.5 ± 1.4 | 93 ± 7 |
| *p* value | 0.9288 | <0.0001 | 0.8987 | 0.7164 | 0.2865 | 0.1043 | 0.2590 | 0.0367 |

Number of neurons n=26 tdTomato- and 23 tdTomato+ for resting membrane potential and input resistance measurements; for the rest of the electrophysiological properties, n=22 tdTomato- and 19 tdTomato+; for the dendritic span n = 12 tdTomato- and 16 tdTomato+.

## ACKNOWLEDGEMENTS

The authors would like to thank members of the Cembrowski and Spruston laboratories, as well as Annie Vogel Ciernia and Jason Snyder, for helpful discussion and comments on the manuscript. Funding was provided by the University of British Columbia (Department of Cellular and Physiological Sciences, Djavad Mowafaghian Centre for Brain Health, and the Faculty of Medicine Research Office), the Natural Sciences and Engineering Resource Council of Canada (RGPIN-2019-04507), and the Howard Hughes Medical Institute.

## METHODS

Experimental procedures were approved by the Institutional Animal Care and Use Committee at the University of British Columbia and the Janelia Research Campus.

### Mouse behavior, activity tagging, and quantification

To label cells active during behavior (Fig. 1), we used a transgenic mouse system that enables permanent tagging of transiently active neurons ("FosTRAP" mice: Guenthner et al., 2013). FosTRAP mice were crossed to Ai14 (tdTomato Cre-reporter) mice (Madisen et al., 2010), which induces tdTomato expression in active neurons following 4-hydroxytamoxifen (4-OHT) administration.

Double-positive mature male mice that received at least three sequential days of transfer and handling in the experiment room were used for behavioral experiments. For novel environment exploration, mice were added to an operant chamber (10.2 in width x 12.6 in length x 8.3 in height) and allowed to explore for 20 minutes. For footstock conditions, mice were placed in an operant chamber and two footshocks were applied (2 s, 0.7 mA; shocks occurring at 2.5 and 3.5 minutes after introduction), and removed after 20 minutes. For novel object experiments, a small plastic toy was added to a mouse’s homecage, and removed after 20 minutes. For novel odor experiments, a cage lid was wiped down with diluted (1:10) peppermint extract, and swapped with the homecage lid for 20 minutes. For experiments involving introduction of a conspecific, male co-housed littermates were individually housed for 3 days, reintroduced into a new homecage for 20 minutes, and subsequently single-housed. For forced swim, mice were placed in a clear acrylic cylinder (12 in height x 10 in diameter) half-filled with warm water and removed after 5 minutes.

Five to fifteen minutes after behavior, 4-hydroxytamoxifen (4-OHT) was administered intraperitoneally to induce tdTomato expression in active neurons (performed as described in Guenthner et al., 2013). After allowing 5-7 days for tdTomato expression, mice were subsequently sacrificed, via deep anesthesia with isoflurane and perfusion with phosphate-buffered saline (PBS) followed by 4% paraformaldehyde (PFA) in 0.1M PB. Brains were dissected and post-fixed in 4% PFA overnight. Brain sections (100 μm) were made using a vibrating tissue slicer (Leica VT 1200S, Leica Microsystems, Wetzlar, Germany). Neurons occupying the granule cell layer or molecular layers, exhibiting a polarized morphology consistent with granule cells, were manually counted around the intermediate dentate gyrus (~- 3.0 mm to bregma). Cells were counted from at least four sections for each animal. At least three animals were used for each behavioral paradigm. Summary statistics are presented as mean ± SEM, with paired Mann-Whitney U tests performed to analyze differences across blades.

To control and help interpret this activity labeling, three sets of negative control experiments were performed (Supplementary Fig. 2A-E). For vehicle control injections, mice received saline rather than 4-OHT following novel environment exposure. For homecage control experiments, mice received 4-OHT within their homecage in their holding room. For behavior room transfer and handle experiments, animals that had received three days of transfer and handling in the experimental room received 4-OHT following a fourth day of transfer and handling in the experimental room.

### Immunohistochemistry

Mature male mice were deeply anesthetized with isoflurane and perfused with 1x PBS followed by 4% PFA in in 0.1M PB. Brains were dissected and post-fixed in 4% PFA overnight. Brain sections (100 μm) were made using a vibrating tissue slicer (Leica VT 1200S, Leica Microsystems, Wetzlar, Germany). Antibodies used in this study were as follows: rabbit antibody to c-Fos (1:500, #2250, Cell Signaling Technology; RRID: AB_2247211), mouse antibody to Prox1 (1:1000, ab92825, Abcam; AB_10563321), rabbit antibody to Pcp4 (1:250, HPA005792, Millipore Sigma; RRID: AB_1855086). Immunohistochemistry was performed on free-floating sections. All tissue was washed 5 times (5 minutes each) in PBS and then incubated in blocking buffer (5% NGS in 0.3% Triton-PBS) for one hour at room temperature. Tissue was subsequently incubated in primary antibody at 4°C overnight, washed 5 times (5 minutes each) in 0.3% Triton-PBS, and detected by Alexa Fluor secondary antibodies (Thermo Scientific Inc., Waltham, MA) by incubating at room temperature for 1-2 hours. Sections were subsequently washed in PBS five times (5 minutes each), mounted, and coverslipped with mounting media containing DAPI (H-1200, Vector Laboratories, Burlingame, CA). Cell bodies were manually counted, as done for quantification of transgenic-tagged cells.

### In situ hybridization

To prepare tissue for single-molecule fluorescent *in situ* hybridization (ISH), mature male mice were deeply anesthetized with isoflurane and perfused with 1x PBS followed by 4% PFA in 0.1M PB. Brains were dissected and post-fixed in 4% PFA for 2-4 hr. Brain sections (20 μm) were made using a cryostat tissue slicer (Leica 3050S, Leica Microsystems, Wetzlar, Germany) and mounted on glass slides. Slides were subsequently stored at −80°C until use. Custom probes for *Arc* (316911-C3), *Cck* (402271), and *Penk* (318761-C2) were ordered from Advanced Cell Diagnostics (ACD, Hayward, CA). Antigen retrieval, pretreatment, hybridization, amplification, and detection were performed according to User Manual for Fixed Frozen Tissue (ACD) (Wang et al., 2012). Cell bodies were manually counted, as done for quantification of transgenic-tagged cells. For two-color quantification of *Cck* and *Penk* expression (Fig. 4H,I), only cells at the GCL-IML border or beyond were examined, as the dense crowding of cells within the GCL precluded segmentation. For comparing activity-labeled *Penk-*expressing cells to that expected by chance, the mean number of tdTomato cells and the mean rate of *Penk* labeling were empirically obtained, and 1,000,000 Monte Carlo simulations were performed where the number of tdTomato-expressing cells were stochastically assigned *Penk* expression based upon chance levels. Statistical significance was assessed by a Mann-Whitney U test, comparing Monte Carlo simulations to empirical values found for 6 animals. Coronal sections from the Allen Mouse Brain Atlas (AMBA) (Lein et al., 2007) were used to perform single-color colorimetric ISH examination of blade-specific differences and scRNA-seq predictions. When quantification was used, cell bodies were manually counted, as done for quantification of transgenic-tagged cells. The genes (experiments) used were *Pcp4* (79912613), *Prox1* (73520980), *Slc17a7* (70436317), *Cck* (77869074), and *Penk* (74881286).

### Surgeries and viral injections

For activation of granule cells, the DREADD virus AAV2-hSyn-DIO-hM3D(Gq)- mCherry (UNC Gene Therapy Center Vector Core) was injected bilaterally into the dentate gyrus in mature Rbp4-cre KL100 male mice (RRID: MMRRC_031125-UCD) (Gerfen et al., 2013) via stereotactic surgery. This line provides selective access to granule cells (Cembrowski et al., 2016b). Injections were located at A/P, M/L, (D/V) −3.3, 2.5, (-4, −3.25, −2.5), and −3.0, 2.0, (-4, −3.25, −2.5), with 80 nL of virus injected at each site. To drive activation, Clozapine N-Oxide (CNO) (BML-NS105, Enzo Life Sciences, Farmingdale, NY, and #4936, Tocris, Bristol, UK) was dissolved in sterile, injectable saline containing 0.5% DMSO. This solution was injected intraperitoneally at 5 mg/kg. For birthdate labeling of granule cells, *in utero* viral injections of AAV2-CAG-FLEX-tdT were performed as described previously (Cembrowski et al., 2016a) into pregnant Rbp4-cre KL100 female mice. Cell bodies were manually counted, as done for quantification of transgenic-tagged cells.

### Mouse hippocampal slice preparation, recording, and morphological analysis

Mature (2-to 4-month-old) FosTRAP x Ai14 mice underwent novel environment exposure and 4-OHT administration, as in behavioral experiments. One week after this, mice were anesthetized with isofluorane and decapitated. The brain was extracted and transferred to ice-cold dissection solution containing (in mM): 80 NaCl, 24 NaHCO_3_, 25 dextrose, 75 sucrose, 2.5 KCl, 1.25 NaH_2_PO_4_, 0.5 CaCl_2_, 5 MgCl_2_, 1 ascorbic acid, 3 Na-pyruvate. The solution was saturated with 95% O_2_ and 5% CO_2_. 300 µm thick coronal hippocampal slices were cut using a vibratome (VT1200S Leica, Germany), then hemisected and placed in artificial cerebral spinal fluid (ACSF) containing (in mM): 126 NaCl, 2.5 KCl, 1.2 MgCl_2_, 2.4 CaCl_2_, 1.2 NaH_2_PO_4_, 11.4 glucose, 21.4 NaHCO_3_, 1 ascorbic acid, and 3 Na-pyruvate saturated with 95%O_2_ and 5% CO_2_ (pH 7.4) and maintained at 32°C. Slices were placed in a submersion-type recording chamber perfused with ACSF at 32°C. Slices were visualized on an upright microscope (BX61WI; Olympus, Tokyo, Japan) equipped with infrared-differential interference contrast optics. The recording pipettes (4–9 MΩ resistance) were filled with internal solution containing (in mM): 130 K-gluconate, 10 KCl, 10 Na_2_-phosphocreatin, 10 HEPES, 4 Mg-ATP, 0.3 Na-GTP, and 0.2% biocytin (pH 7.2, osmolarity 295). Current-clamp experiments in brain slices were performed with whole-cell patch-clamp recordings using a Multiclamp 700B amplifier (Molecular Devices, San Jose, CA). Electrophysiological data were low-pass-filtered with a cut-off frequency of 10 kHz and digitized at 20 kHz via a USB-6343 board (National Instruments, Austin, TX) under the control of WaveSurfer software (https://www.janelia.org/open-science/wavesurfer). Bridge balance and capacitance compensation were performed at the beginning of recordings. After current-clamp recording, cells were kept in whole-cell mode for 30 minutes to allow sufficient biocytin filling of the cell, slices were then fixed in 4% PFA and recorded cells were subsequently detected with an Alexa Fluor 488/streptavidin reaction.

Analysis was performed in Matlab (MathWorks, Natick, MA) with custom scripts. Action potential (AP) amplitude and threshold were calculated as the smallest current in protocol that induced APs (typically 100 – 200 pA). AP was defined as the different in voltage between peak of AP and the resting potential. Adaptation ratio was defined as the interspike interval (ISI) between the first two APs, divided by ISI between the last two APs, within a 2-second 200 pA current injection. Afterhyperpolarization (AHP) was calculated as the difference between the AP trough and the threshold of the AP at 200 pA current injection. Dendritic span was measured as the distance between the two outermost dendrites at 50 µm above the initial bifurcation of primary dendrite, or center of soma if there are multiple primary dendrites in ImageJ (Schindelin et al., 2012). Clustering analysis based on dendritic span and input resistance was done using the k-means clustering algorithm in Matlab. The average percentage of tdTomato positive cells that belongs to a single cluster is 81.3% (averaged value of 1000 calculations using different random number seeds). To verify the robustness of clustering based on dendritic span and input resistance, we performed k-mean clustering for all cells with all electrophysiological properties measurements. The average percentage of tdTomato positive cells that belongs to a single cluster is 94.7% (1000 simulations).

### Single-cell RNA-seq analysis

Computational analysis was performed in R (R Development Core Team, 2008) using a combination of Seurat v1.4.0.16 (Satija et al., 2015) and custom scripts (Cembrowski et al., 2018a; Cembrowski et al., 2018b). Data from a previously published scRNA-seq was used (Habib et al., 2016), with cells annotated as taken from the dentate gyrus used for the analysis here (i.e., those tagged as “DG” from the DATA_MATRIX_LOG_TPM.txt data file). Data were transformed from log to linear space and loaded via *Setup(min.cells=3, min.genes=200, do.logNormalize=T, total.expr=10000).* Subsequent analysis proceeded via default parameters used in the Seurat package. Graph-based clustering was performed on using dimensionally reduced data via principal component analysis, and differential expression between subtypes was assayed via non-parametric Wilcoxon rank sum test. When plotting gene expression in tSNE plots, color ranges from white (zero expression) to red (maximal expression), plotted logarithmically. All analysis scripts will be available upon acceptance or reviewer request.

### Fluorescence Imaging

Images were acquired with a confocal microscope (LSM 880, Carl Zeiss Microscopy, Jena, Germany) using a 20x or 40x objective. Some images were postprocessed in Fiji, including brightness adjustments applied to the entire image, as well as pseudocoloring to facilitate visual comparisons across channels.

