## Supplemental Figures for "A sparse, spatially biased subtype of mature granule cell is preferentially recruited in hippocampal-associated behaviors"

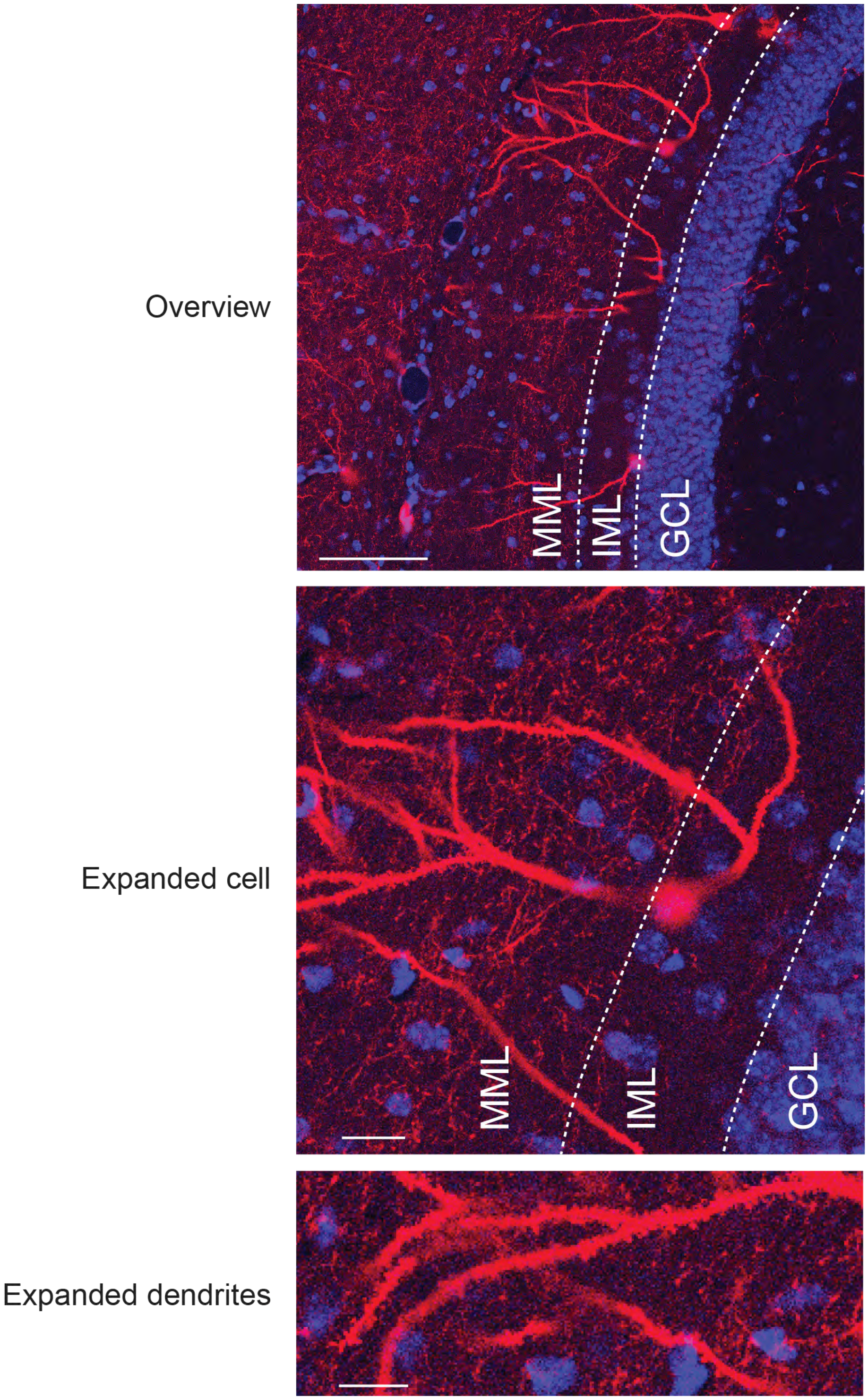


Supplemental Figure S1. Representative depiction of activity-labeled cells and dendrites. Related to Figure 1.
Top: overview of activity-labeled cells in the suprapyramidal blade. Note displaced cell bodies.
Middle: expansion on a labeled cell. Bottom: expansion on labeled dendrites.

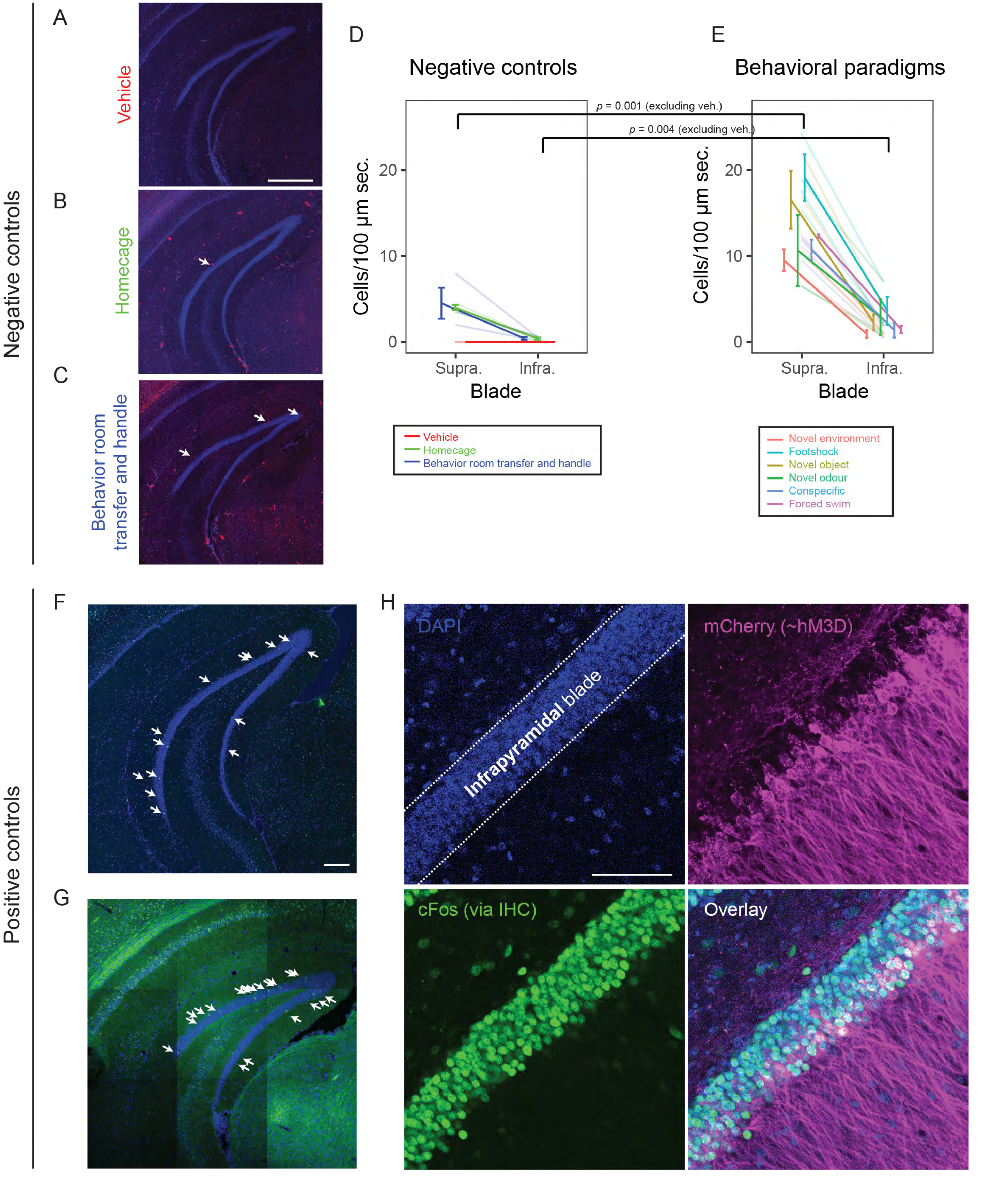


Supplemental Figure 2. Control activity labeling experiments.

A. Representative image showing absence of labeling following vehicle injections. B. Representative image showing activity-tagged cells for an animal in homecage. C. Representative image showing activity-tagged cells for an animal transported to the behavior room and handled, but not put through a behavioral paradigm. Arrows indicate activity-tagged tdTomato-expressing cells in (A-C). D. Summary of negative control data (n=2 mice for vehicle injections, n=3 mice for each of homecage and behavior room transfer). E. For comparison to (D), summary of animals put through behavioural paradigms (as in Fig. 1J). Individual lines reflect individual animals, lines with error bars reflect pooled mean  SEM, and statistical comparisons represent Mann-Whitney U-tests. F. Representative image showing preferential inner blade labeling following novel environment exploration, as assayed through cFos IHC. Arrows indicate cFos+ cells in the granule cell layer and molecular layers. Note in general, the inner blade was associated with 80% of cFos-labeled cells (n = 80/100 cells, 2 animals and 2 sections per animal). G. Representative image showing preferential inner blade labeling following novel environment exploration, as assayed through Arc ISH. Arrows indicate Arc-expressing cells in the granule cell layer and molecular layers. Note that, in general, the inner blade was associated with 70% of Arc-labeled cells (n = 276/395 cells, 2 animals and 6 sections per animal). H. Upper left: DAPI image of the outer blade of the dentate gyrus. Upper right: mCherry expression, corresponding to injection of AAV2-hSyn-DIO-hM3D(Gq)-mCherry into the DG of Rbp4-cre mice. Lower left: immunohistochemical staining of cFos. Similar broad staining of the outer blade of the dentate gyrus was seen in 2 other animals. Lower right: overlay of all panels.

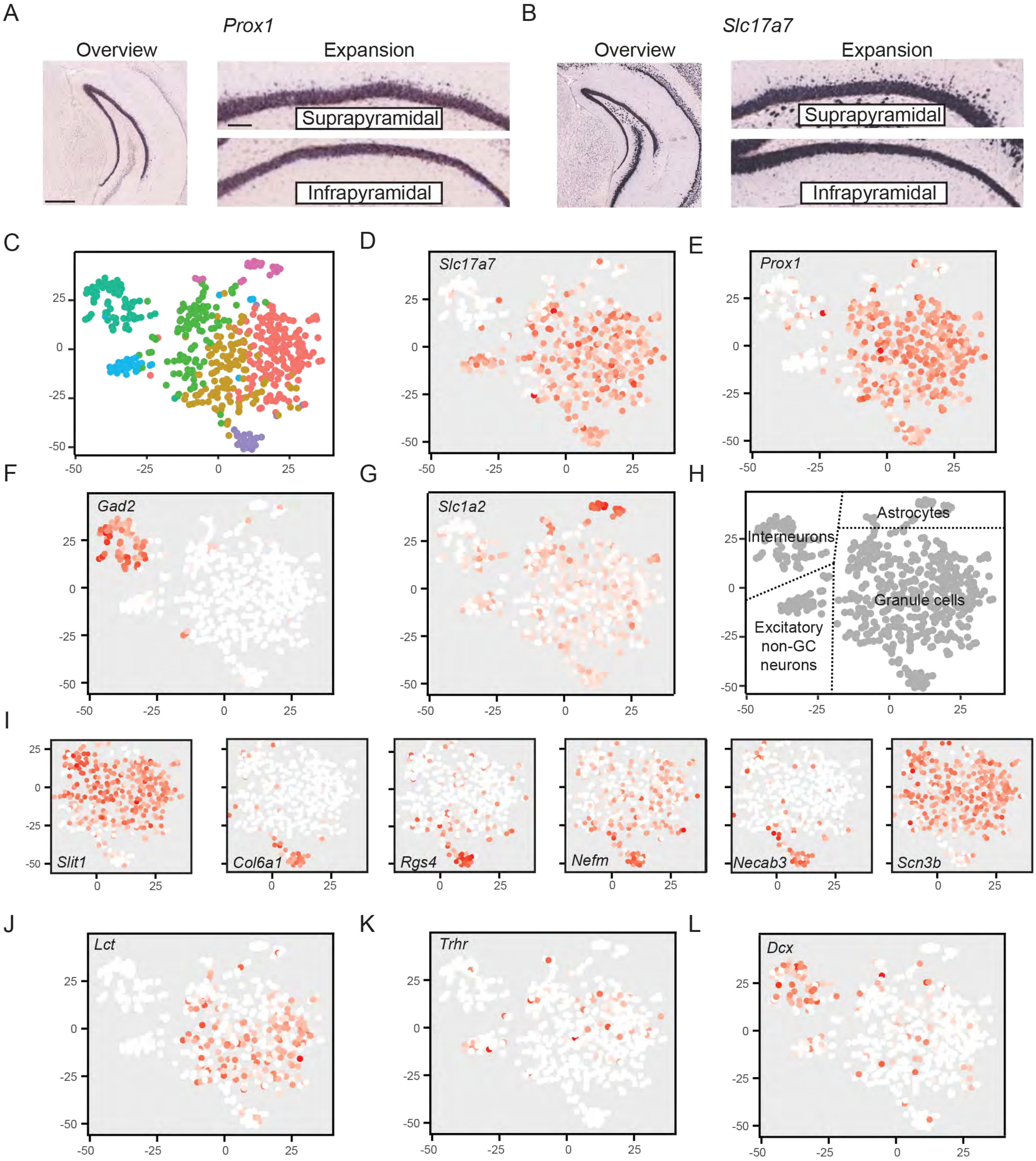


Supplemental Figure 3. Gene expression in the dentate gyrus. Related to Figures 2 and 4.
A. Left: overview of Prox1 expression in the dentate gyrus, a marker of granule cells. Right: expansion of the suprapyramidal (top) and infrapyramidal (bottom) blades. Note suprapyramidal blade enrichment of displaced granule cells. B. As in (A), but for the excitatory neuron marker gene Slc17a7. C. t-SNE visualization of single-cell transcriptomes, with coloring denoting cluster identity. D-G. Expression of the excitatory neuron marker Slc17a7 (D), the DG GC marker Prox1 (E), the interneuron marker Gad2 (F) and the glial marker Slc1a2 (G). H. Cell-type-specific labeling of DG cell transcriptomes. I. Expression of subtype-enriched marker genes with neuronally relevant functional correlates. J,K,L. Expression of the dorsal DG marker gene Lct (J), the ventral marker gene Trh2f2 (K), and the immature granule cell marker Dcx (L). Note that expression of each marker gene in (J,K,L) is dispersed across the GC clusters rather than exhibiting structured expression.
